# A Robust, Low-Cost Instrument for Real-time Colorimetric Isothermal Nucleic Acid Amplification

**DOI:** 10.1101/2021.08.18.456885

**Authors:** F. Myers, B. Moffatt, R. Khaja, T. Chatterjee, G. Marwaha, M. McGee, D. Mitra

## Abstract

The COVID-19 pandemic has highlighted the need for broader access to molecular diagnostics. Colorimetric isothermal nucleic acid amplification assays enable simplified instrumentation over more conventional PCR diagnostic assays and, as such, represent a promising approach for addressing this need. In particular, colorimetric LAMP (loop-mediated isothermal amplification) has received a great deal of interest recently. However, there do not currently exist robust instruments for performing these kinds of assays in high throughput with real-time readout of amplification signals. To address this need, we developed LARI, the LAMP Assay Reader Instrument. We have deployed over 50 LARIs for routine use in R&D and production environments, with over 12,000 assays run to date. In this paper, we present the design and construction of LARI along with thermal, optical, and assay performance characteristics. LARI can be produced for under $1500 and has broad applications in R&D, point-of-care diagnostics, and global health.

## Introduction

Since the invention of the polymerase chain reaction (PCR) in 1985 [1], nucleic acid amplification has become one the most important tools in life science as well as a vital component of many clinical diagnostics workflows. However, one challenge with PCR that has limited its adoption in point-of-care diagnostics settings is the cost and complexity of thermal cycling instrumentation. Benchtop thermal cyclers are bulky and consume a lot of power due to the need for thermoelectric heat pumps, fans, and heat sinks which can deliver rapid active heating and cooling to the assay block. Several isothermal amplification assay strategies have been developed which do not require heat denaturation of DNA to initiate new amplification cycles and thus eliminate the need for temperature cycling. Of these isothermal strategies, Loop-Mediated Isothermal Amplification (LAMP) is perhaps the most well-suited for clinical diagnostic applications [2]. LAMP provides rapid DNA amplification and excellent specificity thanks to its use of four to six primers. Since its initial development in 2000, LAMP has seen many improvements, including the development of reverse transcriptase LAMP (RT-LAMP) for detection of RNA targets [3] new polymerase enzymes specifically engineered for LAMP [4], [5], and lyophilization techniques which enable LAMP reagents to remain stable at room temperature for years [5], [6]. Quantitative RT-LAMP has also been demonstrated based on real-time monitoring of amplification curves [7]. Thanks to its robustness against various inhibitors, RT-LAMP has been demonstrated without nucleic acid enrichment in various clinical specimens as a simplified “single-pot” reaction [8].

Aside from thermal instrumentation, signal readout is another important consideration for simplifying molecular assays. Amplification products are typically detected with fluorescence using either intercalating dyes or molecular probes. This generally requires a high-power light source, optical excitation/emission filters, and a sensitive detector. LAMP generates substantial quantities of DNA (>10^9^ in less than an hour), enabling novel signal readout strategies including turbidimetric monitoring of precipitates [9], electrochemical detection of electroactive DNA intercalators [10], and colorimetric detection [11]–[15]. Colorimetric LAMP offers distinct advantages for readout, as many different indicator dyes are available which produce amplification color signals dramatic enough to be seen with the naked eye. These include metal indicators such as calcein [11] and hydroxynaphtol blue [12], intercalating dyes such as SYBR Green [13], and pH indicators such as phenol red [15], [16].

The simplified instrumentation requirements for colorimetric LAMP make it ideal for point-of-care and resource-limited settings [17], [18]. The LAMP assay can be incubated to a constant temperature with a low-cost battery-powered resistive heating element, and colorimetric readout can be read with a simple LED and photosensor, or even with the naked eye. Colorimetric RT-LAMP assays have been demonstrated for a variety of targets including SARS-CoV-2 [19] and Influenza [20]. Colorimetric LAMP has also enabled molecular diagnostics to enter home use. The Lucira™ All-In-One COVID-19 Test is the world’s first multiplexed nucleic acid amplification platform available for at-home use by consumers [21]. The product uses a pH indicator dye for colorimetric RT-LAMP in a fully-integrated single-use device [22], [23].

To date, colorimetric LAMP has typically been performed with endpoint measurements using imaging solutions for readout such as flatbed scanners [16]. Real-time monitoring of colorimetric signals can provide quantitative information about template concentration [24]. Even when quantitative data is not required, real-time color change data is nevertheless very useful. For example, it can reduce assay result times, as signal slope changes can often be detected sooner than absolute color shifts. Real-time monitoring also allows time cutoffs to be applied to amplification times to manage risks of false positives (e.g. from non-specific amplification) and target a particular sensitivity/specificity point on the assay’s receiver-operator curve (ROC). This is particularly important with isothermal assays, which can continue amplifying and changing colors even after an assay’s designated runtime.

There is not currently commercial instrumentation specifically designed for real-time monitoring of colorimetric amplification assays. While conventional qPCR thermocyclers can be used for isothermal amplification, their optical systems are configured for epifluorescence and generally cannot monitor optical absorbance of light through a sample. This makes them unsuitable for real-time monitoring of colorimetric LAMP signals. Plate spectrophotometers, on the other hand, can be used for real-time monitoring of absorbance through a specimen and some of these provide incubation functions. One colorimetric LAMP assay for SARS-CoV-2 was recently granted emergency use authorization using such an instrument [25]. However, sample plates are heated via convection, which increases assay time and raises the risk of nonspecific amplification at intermediate temperatures as the assay warms up. Given these limitations, there is a need for new instrumentation specifically tailored for colorimetric isothermal amplification. Such instrumentation has broad applications in R&D, clinical diagnostics, global health in resource-poor settings, and in-field analysis for agriculture or environmental applications.

In this paper, we present the LAMP Assay Reader Instrument, or LARI, an instrument designed for multiplexed isothermal nucleic acid amplification with real-time colorimetric readout. We developed LARI to conduct R&D and quality control monitoring of colorimetric LAMP assays in a production environment. To date, over 50 LARI instruments have been built and deployed across multiple sites, and these instruments are used on a daily basis.

This small benchtop instrument functions much like a conventional qPCR thermal cycler and includes a heating block for up to 13 0.2 mL PCR sample tubes. Unlike a PCR thermal cycler, however, the instrument does not include active cooling and is only designed to heat samples to a constant temperature. It also features LEDs and photosensors which monitor changes in light transmission through sample vials. This paper presents the design and construction of LARI and evaluates the instrument’s thermal, optical, and colorimetric LAMP assay performance.

## Materials and Methods

### Instrument Design

Figure 1 shows a cross-sectional view and photographs of LARI. Table 1 lists the critical components. LARI accepts a standard strip of 0.2 mL PCR sample tubes. Twelve tubes locations are available for use, with a 13^th^ tube location included for self-referencing and calibration (currently unused). During use, the lid is opened and tubes are inserted into a 1” x 1.03” x 6” machined aluminum heating block. A second heating block in the lid (1” x 0.75” x 6”) rests on the tops of the tubes when the lid is closed. This lid heating block is held at a higher temperature than the tube heating block to prevent condensation on the inside of the tube lids during a run. It contacts the tube tops via a piece of high-temperature rubber. Both heating blocks are heated via 100W cartridge heaters which are press-fit into through-holes in the blocks with a thermal compound. The tube block includes two cartridge heaters on opposite sides and the lid block includes one in the center. Both blocks include a 100 Ω bolt-on resistive thermal device (RTD) which provide temperature feedback for thermal regulation.

**Figure 1.**
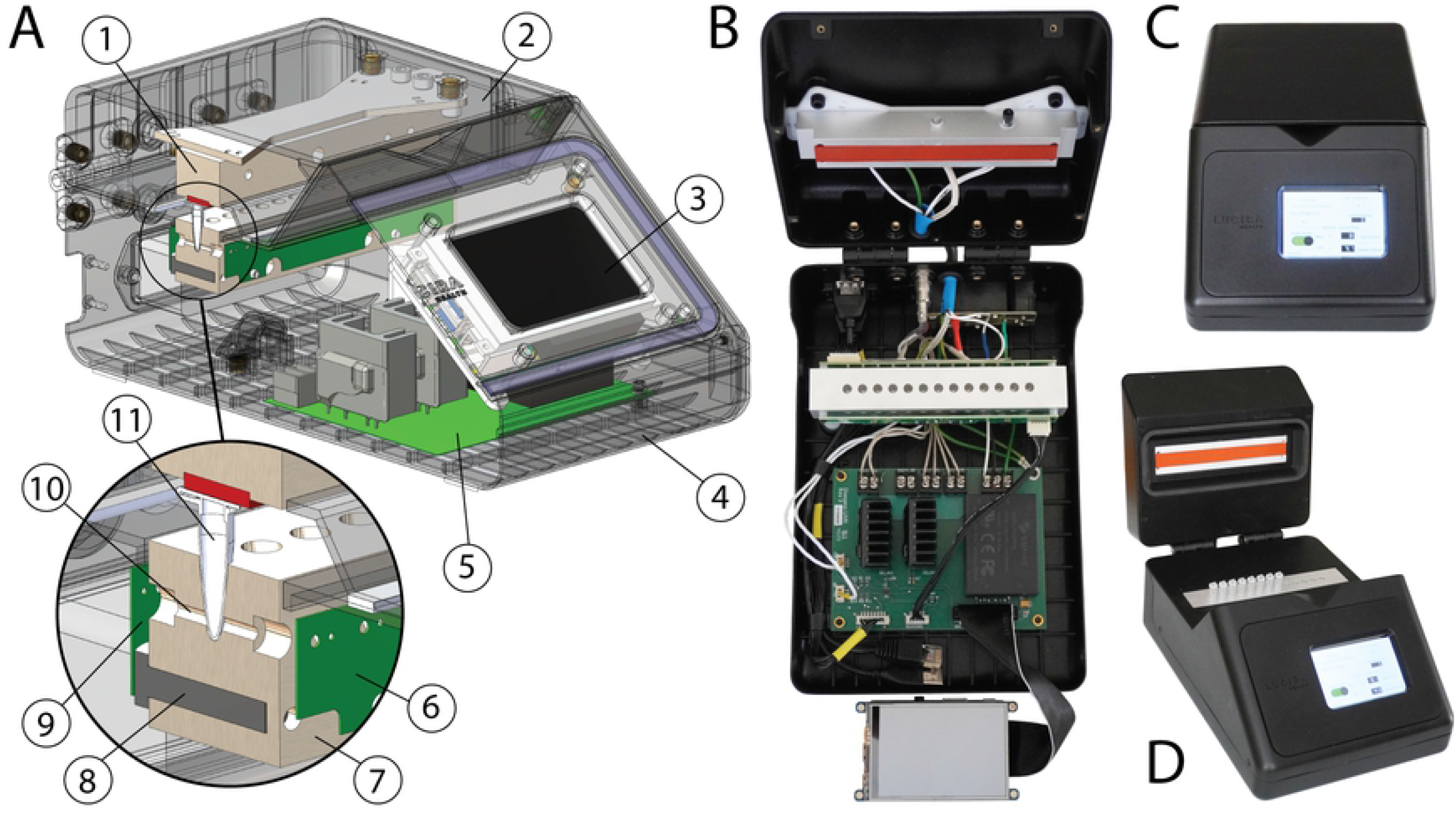
(A) CAD cross section of LARI: (1) A Lid heating block, (2) Lid housing, which is hinged to the base housing, (3) Raspberry Pi single-board computer with LCD touchscreen interface, (4) Base housing, (5) Main control PCB, (6) Sensor PCB, (7) Tube heating block, shown in cross-section through a tube pocket and cartridge heater, (8) Cartridge heater, (9) LED PCB, (10) Through-holes in heating block allow light transmission from the LEDs, through the sample tubes, and on to the sensors, (11) Sample tubes are loaded into machined pockets in the block. (B) LARI shown with the housing open. All components are visible including the lid and tube heating blocks, PCBs, Raspberry Pi/touchscreen, and cabling. (C) Fully assembled LARI with lid closed. (D) Fully assembled LARI with lid open, showing sample tubes loaded.

**Table 1:** Bill of materials for critical components

| Component | Mfg | P/N | Qty |
| --- | --- | --- | --- |
| *LED Board* |  |  |  |
| 590 nm Yellow LED | BiVar | SM1206NYC-IL | 13 |
| LED Driver | Texas Instruments | TLC5928PWPR | 1 |
| *Sensor Board* |  |  |  |
| Digital Light Sensor | AMS | TCS34717FN | 13 |
| I2C Bus Switch | Texas Instruments | PCA9548ADBG4 | 2 |
| *Main Board* |  |  |  |
| ADC | Texas Instruments | ADS1015IDGSR | 1 |
| Solid State Relay | Crydom | PF240D25 | 2 |
| 5V Power Supply | CUI | PSK-20B-S5 | 1 |
| *Computer* |  |  |  |
| Single Board PC | Raspberry Pi | Model B | 1 |
| LCD Touchscreen | Adafruit | 2097 | 1 |
| *Heating Blocks* |  |  |  |
| Cartridge Heater | Heating Elements Plus | TD25010BR | 3 |
| RTD | Omega | RTD-850-B | 3 |
| TCO | Cantherm | L5011024DELB0XE | 2 |
| Thermal Paste | Omega | OT-201-2 | N/A |

Two PCBs on either side of the tube heating block provide optical absorbance readings: one PCB contains LEDs and the other PCB contains digital light sensors. LEDs and sensors are individually addressable, and each tube location has a dedicated LED/sensor pair. Through-holes in the block allow light to transmit from the LED, through the sample tube, and onto the light sensor. The effective path length through the tube is 1.2 mm, assuming a typical 0.2 mm tube wall thickness. These PCBs are separated 3.2 mm from the heating block with standoffs to avoid excessive heating of the electronic components.

The instrument is housed in a 5-piece custom urethane cast housing (Hei-Cast PU8150) which is fastened together with threaded inserts and stainless-steel screws. Small magnets in the lid provide slight downward pressure on the tubes when the lid is closed. The heating blocks are positioned off the housing using nylon standoffs to provide thermal isolation.

Figure 2 shows an overall block diagram of the system. Control software runs on a Raspberry Pi Model B with an LCD touchscreen user interface (UI). The UI allows the user to adjust runtime and setpoint temperature, initiate pre-heating, and initiate or abort an assay run. The Raspberry Pi uploads data to a cloud file server hosted by Egnyte. The main PCB includes two solid state relays which deliver power to the cartridge heaters in the heating blocks. The main PCB also includes a two-channel, 12-bit analog-to-digital converter (ADC) which reads differential voltages from two 100-ohm Wheatstone bridges connected to the RTDs. The control software regulates temperature in the heating blocks using a proportional-integral-differential (PID) control algorithm which monitors temperature feedback from the RTDs and modulates heater PWM duty cycle accordingly. The Raspberry Pi and other electronics are powered by a 5V AC-DC power supply on the main PCB.

**Figure 2.**
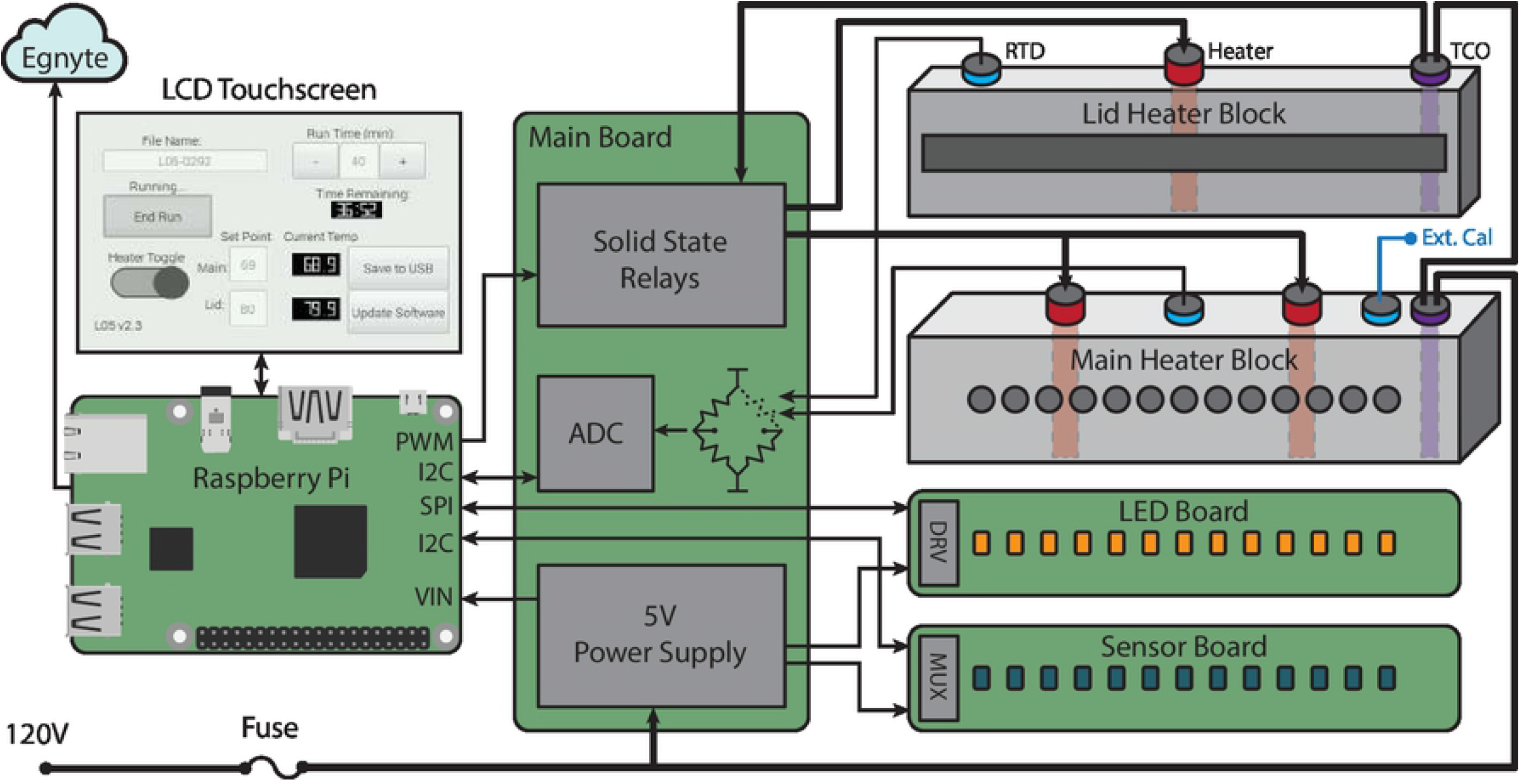
LARI block diagram, including all principal components and touchscreen UI layout.

LEDs were chosen for their brightness (1000 mcd), narrow viewing angle (30 deg), and with a peak wavelength that aligns with the primary absorption wavelength of Lucira’s pH indicator dye (590 nm). The LEDs are driven by a 16-channel constant-current LED driver which communicates to the Raspberry Pi via a Serial Peripheral Interface (SPI). The sensors communicate to the Raspberry Pi over an Inter-Integrated Circuit (I2C) bus through a multiplexing switch IC on the sensor board. During an assay run, the software sequentially illuminates each LED and records the light intensity reading on the corresponding light sensor.

The instrument runs on 120V AC. Input power is routed through a 5A fast-blow overcurrent protection fuse as well as two thermal cutoff (TCO) fuses, one in each heating block. These ensure that power is removed from the device if either block exceeds 110° C.

### Software and User Workflow

Control software is implemented in Python with a QT graphical touchscreen user interface. An overall user workflow is provided in Figure 3. At power-up, the software performs a self-check of all LEDs and sensors to confirm the on/off values of the LEDs are within expected ranges. Prior to starting an assay run, the user is able to adjust the setpoint temperature and runtime for an experiment. Once the user presses the “Pre-Heat” button, the lid block is heated to 80° C and the tube block is heated to 30° C. This ensures a consistent starting temperature across assay runs, regardless of ambient temperature. Once both blocks reach their setpoint temperatures, the user inserts the sample tubes and starts the run. At this point, the tube block will heat to its designated setpoint with a controlled heating ramp profile which may include a short-term hold at a lower temperature. This is helpful for assays involving a reverse transcriptase (RT) enzyme which runs optimally at a lower temperature than the polymerase. The heating ramp profile is software-defined. During a run, the software spawns three execution threads. The first thread cycles each LED and records corresponding sensor values. Each tube is sampled every 10 s. The second and third threads run a PID control loop for the heaters at a 3 s loop interval. Once the specified run duration has elapsed, the heaters are turned off and data from the run is uploaded to an Egnyte cloud file storage database as a comma-separated value (CSV) file with a unique assay ID. The user then removes the sample vials. Before starting a new run, the instrument is allowed to go through a cooldown cycle back to its pre-heat temperature. Although the instrument does not include active cooling, cold blocks can be used to speed up the cooling process and prepare for the next run.

**Figure 3.**
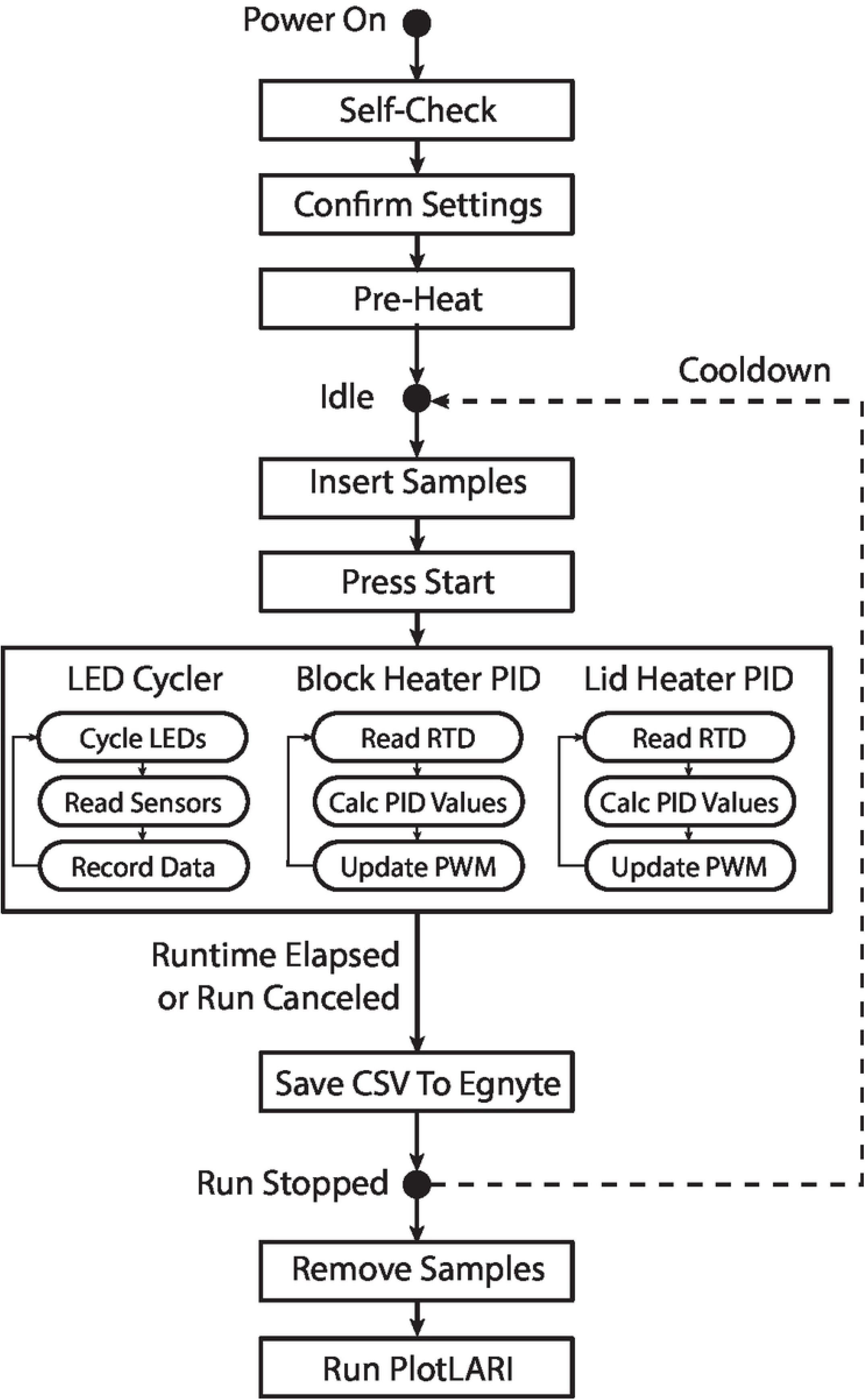
LARI user workflow.

After finishing a run, the user runs a Matlab analysis script on a PC which (a) plots raw data, (b) plots normalized data, where each signal is normalized against its corresponding value at t = 5 min, and (c) calculates time-to-result (TTR) values for each tube in the device. TTR is analogous to the Ct value in PCR and corresponds to the time at which amplification is first detected. In this study, we define TTR as the time at which signal amplitude increases by 50% of its baseline amplitude at t = 5 min. If this condition is met, a tube is considered positive and a TTR value is reported. If this condition is not met, the tube is considered negative and a TTR value is not reported.

### Thermal Performance Evaluation

To characterize transient temperature regulation, miniature K-type thermocouples (Omega SCASS-020G-6-SHX) were inserted into 30 μL of DI water in sample tubes which were then loaded into the instrument at various tube positions. A datalogger (Omega RDSL65SD-USB) recorded the temperature across each tube as the instrument ramped from its pre-heat temperature of 30° C, through a 1 min hold cycle at 50° C, and to a final setpoint of 69° C. Steady-state temperatures were measured for all tubes across five LARIs where a single thermocouple was inserted into each tube one-by-one and allowed to reach equilibrium.

Thermal images were recorded at steady-state with a Fluke TiS45 infrared camera with electrical tape covering the heating blocks for emissivity matching.

### Optical Performance Evaluation

Optical sensitivity to absorbance of reaction solutions was characterized using Black PN dye (TCI B1561, MW = 867.66 g/mol). This dye was chosen for its strong absorption peak at 590 nm and its relative stability with respect to temperature and pH. A stock solution was prepared at 5 mg/mL (5.76 mM) and serially diluted 1:2 in DI water. Absorbance at 590 nm was measured for each dilution in a cuvette using a BioTek Epoch 2 spectrophotometer. To measure corresponding signal amplitudes in LARI, 100 μL of dye at each concentration were aliquoted into a strip of 12 optically-clear PCR tubes (VWR 93001-118). The instrument was allowed to reach a steady-state temperature of 69° C before the first strip of tubes was loaded. After 10 minutes, the tubes were swapped for the next concentration in the series. Finally, the mean and standard deviation values were calculated for each concentration step. The instrument response curve was fitted to a 4-parameter logistic regression (4PL) model. Inflection point, slope, and signal at inflection point were then extracted from this model and compared across tube locations and across 5 LARIs. Coefficient of variation (CV) was estimated across all concentrations. Detection limits were defined as the range of concentrations corresponding to a signal-to-noise ratio (SNR) >2. In other words, minimum detectable concentration was estimated as 2CV above the maximum of the 4PL curve; maximum detectable concentration was estimated as 2CV below the minimum of the 4PL curve.

### Colorimetric LAMP Assay Evaluation

To characterize LARI’s performance on colorimetric LAMP assay performance, signal readout and time-to-result (TTR) performance were evaluated across multiple instruments as well as across tube locations within an instrument. Two different assays known to produce consistent signal and TTRs were used for instrument characterization. Each assay was lyophilized into a room-temperature stable pellet which was dispensed into 0.2 mL PCR tubes. Assay pellets contained DNA polymerase and LAMP primers. One pellet contained a co-lyophilized target (positive control) and one did not (non-template control, NTC). Pellets were hydrated in a weakly-buffered solution containing a pH indicator dye with a peak absorption wavelength at 590 nm. During DNA polymerization, the solution acidifies due to dNTP incorporation and causes the indicator dye to change color from a dark purple to light yellow (see Figure 6).

To assess instrument-to-instrument assay variability, a single assay manufacturing lot was run across five different LARIs, with six runs per LARI. Each run included four replicates of positive control assay (loaded in tube locations T4-T8), for a total of n=120 assay amplification signals. The resulting signals were analyzed for pre-amplification signal amplitude at t = 5 minutes and post-amplification signal at t = 15 min. TTR was calculated as well as the slope of signal change at TTR and the ratio of post-to pre-amplification signal amplitude. Variation across instruments was analyzed using one-way ANOVA for all parameters to determine if instrument construction variability affects assay performance.

To assess tube-to-tube assay variability, positive control and negative control assays were run across all tube locations. (n=30 runs per assay, with 10 manufacturing assay lots and 15 LARIs represented). The same parameters were calculated as for instrument-to-instrument variability. Variation across tube locations was analyzed using one-way ANOVA for all parameters to determine if tube location affects assay performance.

## Results

### Thermal Performance

Figure 4A shows the transient temperature response of LARI. Tube liquid temperatures lag the block temperature by approximately 20 s during ramp up from room temperature. This lag is more pronounced for tubes at the edge of the block (Tube 1, Tube 4) than tubes at the center (Tube 7). During temperature ramp, LARI draws a peak mains current of about 2.4A. At steady state, tube temperature is about 1° C cooler than the block temperature, as shown with the calibration RTD probe. Software-reported block temperature is calibrated to match closely with tube temperature at steady state. Figure 4B shows a thermal image of LARI with the lid open, with good uniformity across the heating block and lid block. When closed, the lid is the hottest portion of the enclosure, with a maximum surface temperature of 45.6° C (assuming a steady-state block temperature of 69° C and lid temperature of 80° C). Figure 4C shows steady-state temperature uniformity within sample tubes. Steady-state temperature at a 69° C set point varies across all tube locations within an instrument by no more than ±0.4° C. Average temperatures across five LARIs vary by ±0.2° C.

**Figure 4.**
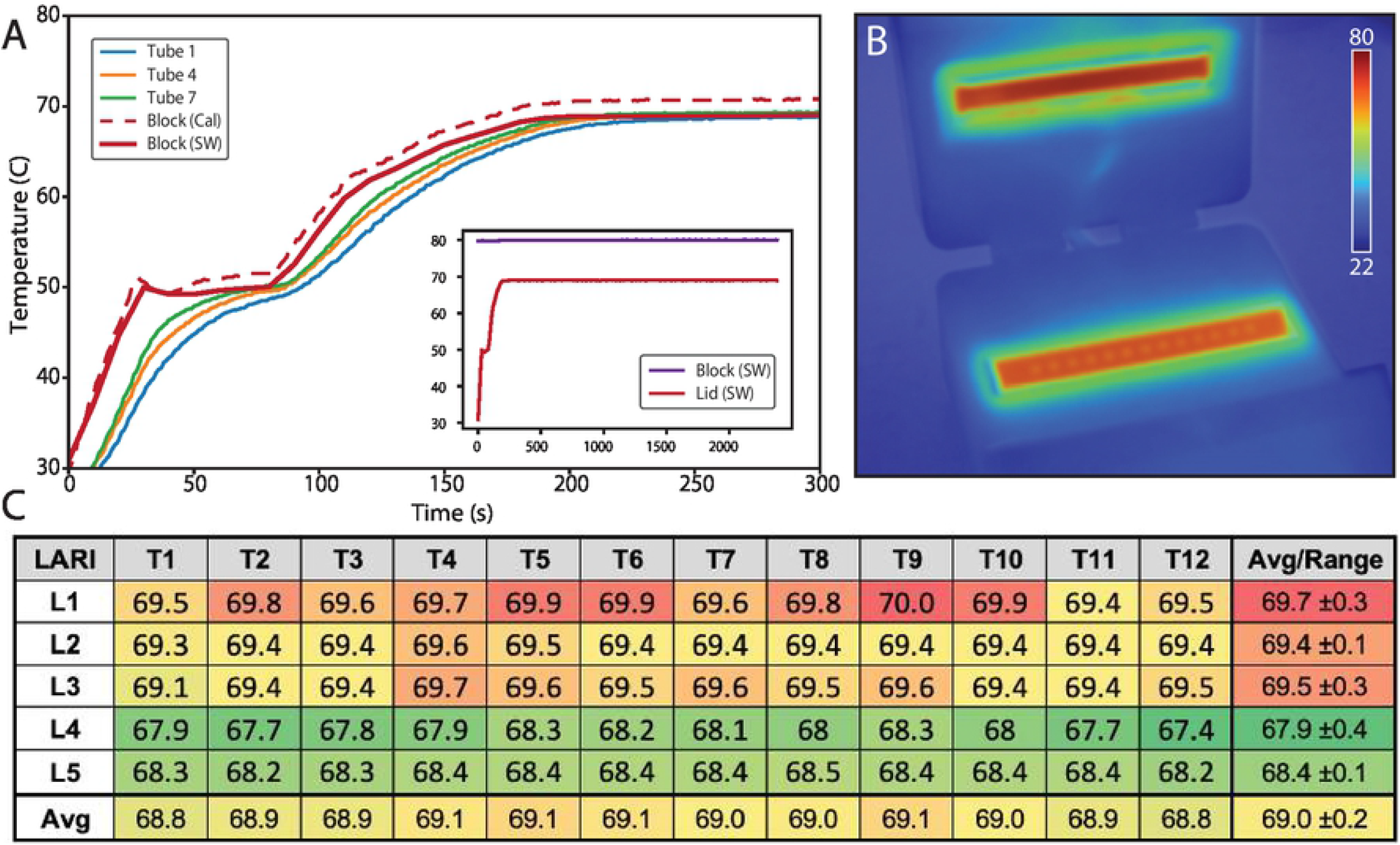
Thermal performance of LARI. (A) Transient temperature response over two temperature set points as measured at three different tube locations: T1 (leftmost tube), T4 (directly over left cartridge heater), and T7 (center of block). Also shown are the software-reported temperature readings as well as temperature monitored with the calibration RTD within the block. Tube temperature lags block temperature by about 10-20 s, with outermost tube lagging the worst. Inset shows full 40 minute run. (B) Steady-state thermal camera image of LARI with tube block at 69° C and lid block at 80° C. (C) Steady-state sample tube temperature across 5 LARIs as measured with an external thermocouple inserted in water in each tube, showing excellent temperature uniformity.

### Optical Performance

PN dye spectrum and absorbance at 590 nm as a function of concentration are provided in the supplemental information (see S1). Absorbance units are normalized to a 1 cm path length. A linear regression (with y-intercept of 0) was fit to this curve to obtain an extinction coefficient, ε = 30,289 Lmol^-1^cm^-1^.

Figure 5A shows an example LARI run with successive dilutions of PN dye introduced into all 12 tube locations at five minute intervals. Signal intensities from the light sensors are provided as digital counts (cnts). At t = 0 min, the instrument has no tubes installed and the signal saturates (s = 2^16^ cnts). At t = 5 min, tubes containing DI water are introduced. At t = 10 min to t = 75 min, serial 1:2 dilutions are successively introduced, starting with 5762.6 µM (5 mg/mL). At t = 75 min, DI water is again introduced. Figure 5B shows the aggregate results of this experiment across five LARIs for dye concentrations within the instrument’s most sensitive range. The instrument’s response follows a sigmoidal curve within this concentration range. Fitting this curve to a four-parameter logistic regression (4PL) yields a midpoint of 223.6 µM, which corresponds to 6.78 absorbance units (Au) at 590 nm. This is the range of maximum optical sensitivity for LARI, where signal changes at a rate of -9.45 cnts/µM, or -311.9 cnts/Au. Signal variability across the full concentration range assessed was very consistent, with an average coefficient of variation (CV) of 37.6%. The minimum and maximum signal readings estimated from the 4PL model were 11.9 and 28,469.4, respectively. Based on this signal range and a SNR requirement of >2, detection limits were determined to be 85.8 — 1274.5 mM of PN dye, or 2.6 – 38.6 Au at 590 nm.

**Figure 5.**
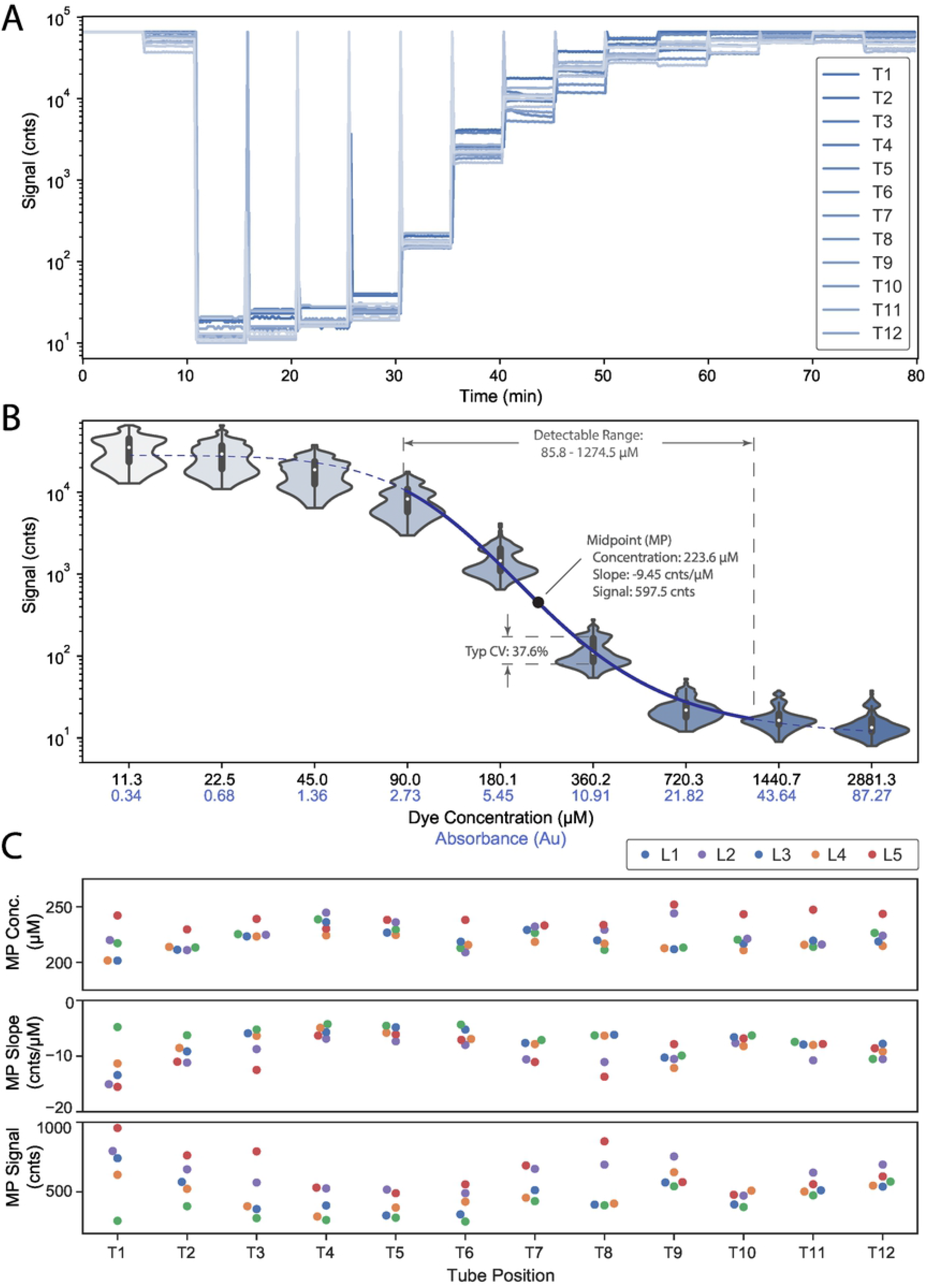
Characterizing LARI signal performance using black PN dye. (A) Signal plots across all 12 tubes in a single LARI run where a serial dilution series of dye samples were swapped in at 5 min intervals. (B) Average signal readings vs. dye concentration across 5 LARIs (12 tubes per LARI). These readings were fit to a 4-parameter logistic (4PL) regression model as shown with the dotted line. (C) 4PL parameters (midpoint, slope, and signal amplitude) do not show significant differences among different tube positions or LARIs, indicating variability is primarily due to LED and sensor variability.

Figure 5C shows 4PL instrument-response parameters for each LARI instrument and tube location evaluated. One-way ANOVA indicates statistically significant differences in these parameters (midpoint concentration, slope, and signal), with the most pronounced variability occurring across LARI instruments (see S2). We attribute this variability primarily to tolerance stack-up of LED brightness, LED drive electronics, and sensor sensitivity. A weak trend in midpoint signal brightness is also observed across tube locations, with tubes at the edges of the block producing brighter signals than tubes at the center of the block. This may be due to slightly higher temperatures at the center of the block attenuating the LED brightness in those locations.

### Colorimetric LAMP Assay Performance

Figure 6A shows typical signal curves from a particular colorimetric LAMP control assay run. Signals start at a mean of 721 counts at t = 5 min for both positive and negative signals; positive signals reach a mean of 3875 by t = 15 min, a ratio of 5.37x. Signals continue climbing beyond 15 minutes and begin to plateau between 4000 – 6000 counts by t = 20 min. Although curves show significant amplitude variability from tube-to-tube (CV at 5 min and 15 min for positive curves is 14.7% and 13.4%, respectively), amplification timing is consistently detected, with TTR values tightly clustered around a mean of 9.18 min (CV = 4.0%) in this particular example. This is because TTR is based on relative changes in signal (normalized to t = 5 min), rather than absolute changes. The inset photograph is a representative example of dye color for positive and negative results.

**Figure 6.**
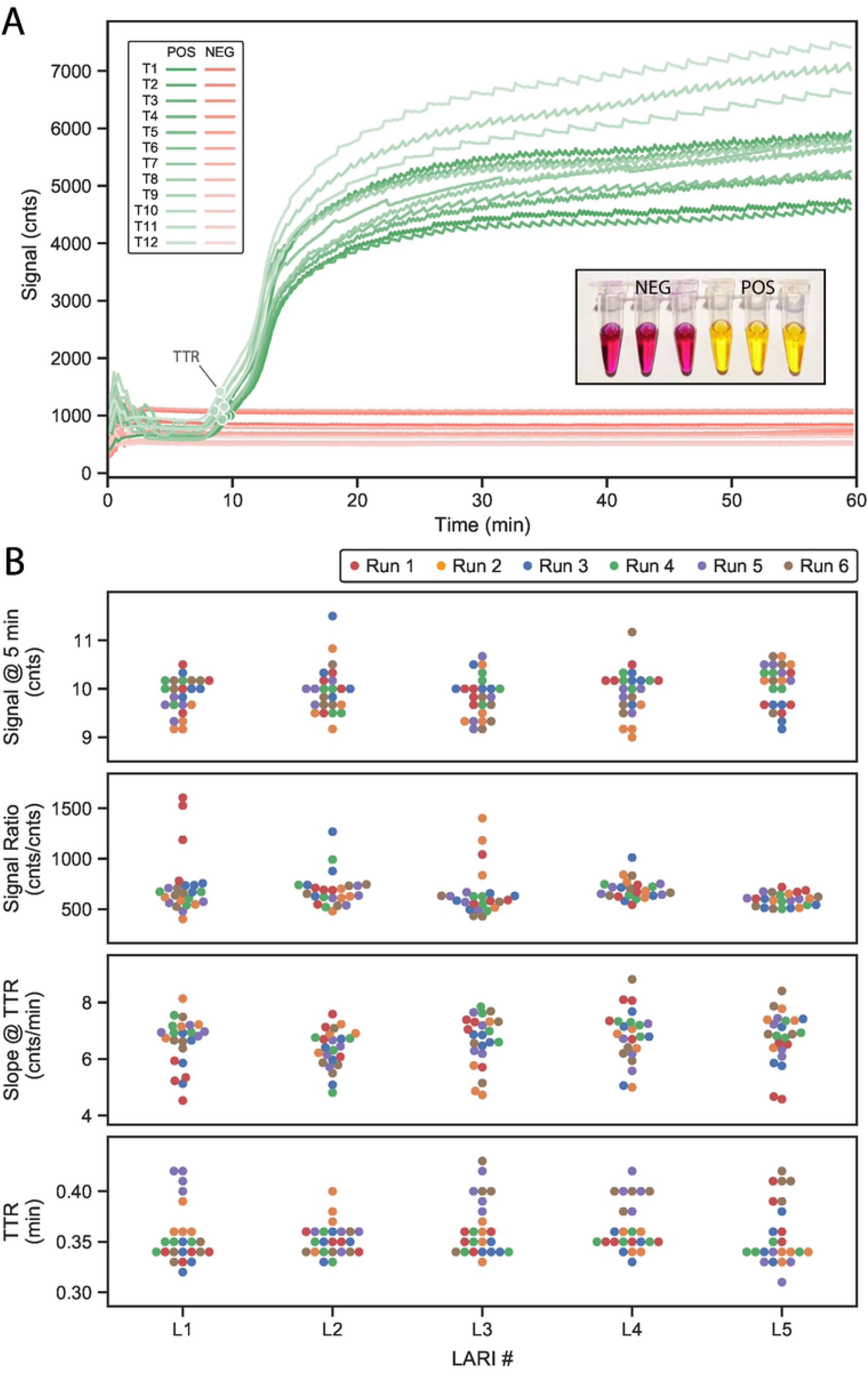
(A) Example colorimetric LAMP assay signals across all 12 tube locations from two different assay runs: one with all positive samples and another with all negative (NTC) samples. Markers indicate TTR for each curve. Inset image shows buffer color at the end of an assay run for both positive and negative samples. (B) Colorimetric LAMP assay performance across six independent runs across five LARI instruments, all using the same manufacturing lot of assay reagents. No discernable trends are visible either from run-to-run or instrument-to-instrument.

Figure 6B extends this experiment across multiple LARI instruments, all using the same lot of positive control assay reagents. A one-way ANOVA did not indicate any significant differences in TTR, starting signal amplitude (t = 5 min), amplification signal ratio (15 min / 5 min), or signal slope at TTR (see S3). Mean TTR value was 9.93 min with a CV of 4.5%.

Figure 7 shows results of positive and negative assay runs across all 12 tube locations. A one-way ANOVA did not indicate significant differences in signal amplitudes, signal ratio, or TTR slope across tube locations (see S4). Statistical differences were observed in TTR, with tubes in the middle of the heating block producing slightly faster TTRs than tubes at the edges. The largest TTR difference observed was 19 s in the positive control assay between T1 (left edge) and T9 (middle). This is attributed to the aforementioned temperature lag across the heating block; the middle tubes heat up sooner and therefore have slightly faster amplification times. This experiment includes 10 different lots of assay reagents. Mean TTR value was 9.49 min with a CV of 3.8%.

**Figure 7.**
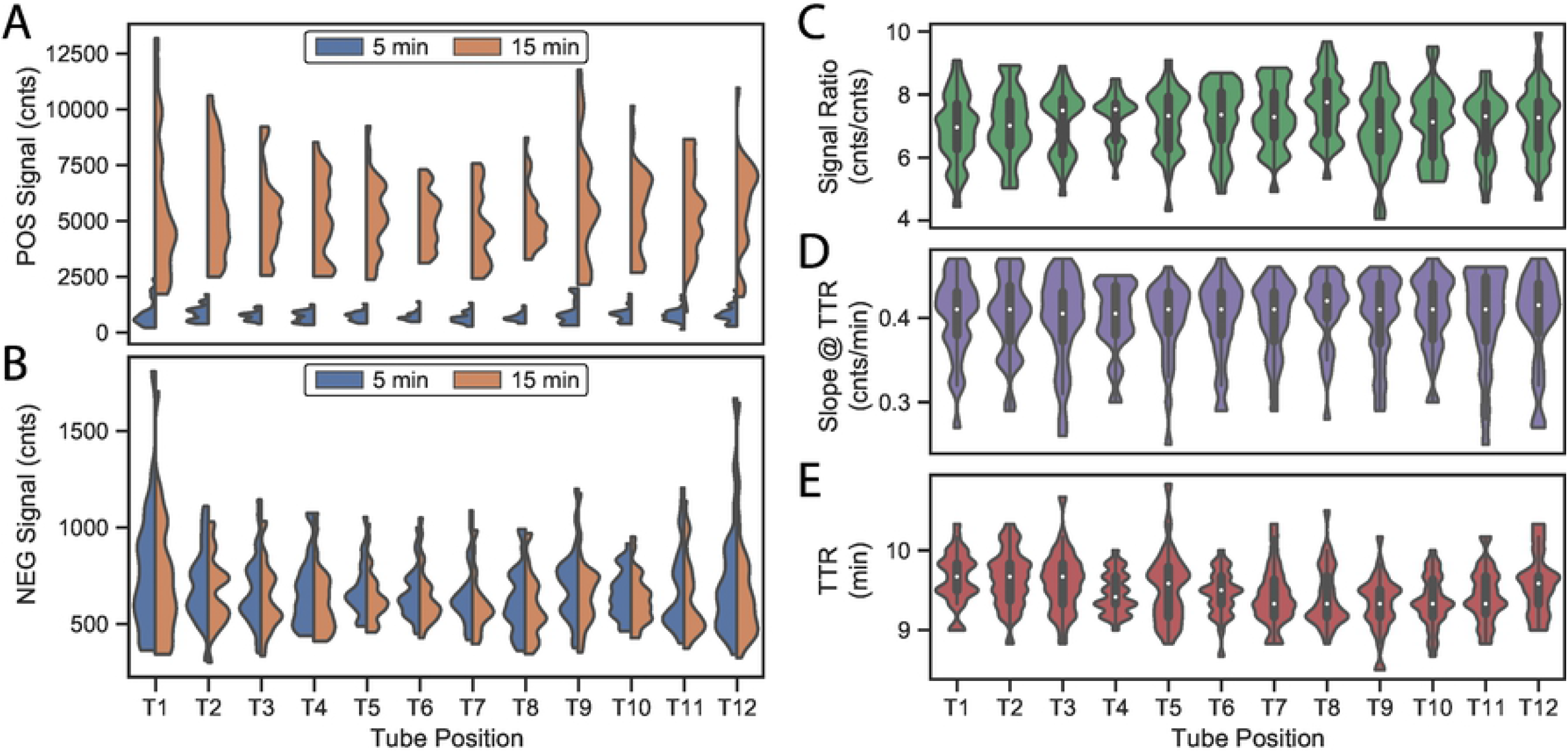
(A) Signal amplitudes for positive assays across all tube positions at t = 5 min (before amplification) and t = 15 min (after amplification). (B) Signal amplitudes for a negative assay at the same timepoints. (C) Ratios of signals for positive assays at t = 15 min to signals at t = 5 min all tube positions. (D) Signal slope at TTR for positive assays across all tube positions. (E) TTR for positive assays across all tube positions.

## Discussion and Conclusion

The LARI instrument presented here can be built for under $1500 in small quantities, including custom PCB assembly, low-volume urethane casting of housing components, and custom machining of heating blocks. As such, it represents a simple, affordable platform for performing multiplexed colorimetric LAMP assays with real-time readout in a convenient benchtop form factor using standard 0.2 mL PCR tubes.

Instrument-to-instrument variability did not appear to meaningfully contribute to assay result variability, indicating that the design and construction methods are robust. This has allowed us to scale up LARI to support our assay QC operations with the same acceptance criteria applied across 50 LARI instruments. Before being placed into operation, each LARI undergoes an installation qualification, operational qualification, and performance qualification (IQ/OQ/PQ) procedure in which, among other things, temperature and signal performance are evaluated to ensure they are within normal operating ranges. LARIs are re-qualified at a 6-month interval.

Instrument performance was also evaluated to determine if tube location contributes to assay result variability. We did find that tubes toward the center of the heating block heated slightly faster and produced slightly faster amplification times, with a difference of 19 s between the fastest and slowest tube locations. This variability is acceptable for our purposes. If better heating and TTR uniformity across the heating block is required, the heating block could be enlarged to reduce thermal edge effects and reduce the variability in temperature ramp profiles across the block. Steady-state temperature, however, showed excellent uniformity on par with conventional PCR thermocyclers.

LARI’s instrument response curve is highly non-linear and signal amplitude is highly variable. As such, signal amplitude is not meant to be interpreted quantitatively. Rather, the instrument is designed to quantify timing of signal changes (i.e. TTR). The instrument is sensitive to dyes absorbing at 590 nm with a dynamic range of 2.6 – 38.6 Au. Dye concentrations outside of this range cannot reliably be distinguished from blank or saturation (i.e. min or max) readings, given measurement noise and instrument variability. One reason for this limited dynamic range is the use of conventional PCR strip tubes. Refraction around the curved walls of these tubes contributes to a background signal even at high (opaque) dye concentrations. Likewise, the limited 1.2 mm path length through the bottom of these tubes reduces sensitivity at low dye concentrations. A wider dynamic range could be achieved using a custom consumable with flat sidewalls and a longer path length—similar to a conventional cuvette. To further reduce background light from stray reflections, the aluminum block could be anodized black.

In our testing with PN dye, we observed absolute signal variability of 37.6%, and this was consistent across a wide range of dye concentrations. Signal amplitude variability can be attributed primarily to variability in the electronics, including the LEDs, LED drive electronics, and sensors. Of these, LED brightness variability is likely the biggest contributor of signal variability. Statistical differences were observed in 4PL instrument-response parameters across tube locations and across instruments when evaluated with a common set of serially-diluted PN dye samples. However, similar statistical differences were not observed in assay starting signals, signal ratios, or signal slopes when using actual replicates of a colorimetric LAMP assay, suggesting that variability of the assay itself (including starting pH, dye concentration, and other sources of experimental variability) contributed more variability than the instrument did in these experiments. To reduce signal amplitude variability, LEDs could be binned from the vendor or during assembly using an optical power meter. Alternatively, instruments could undergo a calibration routine whereby signals are recorded and a lookup table is programmed into each unit to normalize sensor values against a known calibration standard. This type of calibration is commonly performed on other sensitive optical equipment.

The positive control assay presented here includes co-lyophilized template nucleic acid at moderate concentration within the assay pellet itself. As such, this particular assay provides the most repeatable TTR of any assay we test and is a good tool for benchmarking instrument performance. Calculated TTR values showed a CV of 4.5% for a single assay lot run across multiple experiments and multiple instruments. When expanded to multiple lots of assay reagents, TTR CV was similar, at 3.8%. The ten lots used in this experiment were manufactured over a period of several months. This suggests that lot-to-lot variability of assay reagents is not a significant contributor to the TTR variability observed on LARI. If we assume the observed TTR variability is entirely due to the instrument itself, we can conclude that LARI is capable of reporting TTR values of colorimetric assays with a precision of 4.5% or better.

To date, we have deployed over 50 LARI instruments across QC, R&D, and clinical collaborators. Collectively, these instruments have completed over 12,000 isothermal colorimetric LAMP assay runs. The instruments have run reliably with relatively few problems. Our main challenge has been drift in LED brightness, resulting in instruments occasionally failing to meet signal specifications during periodic re-qualification testing. When this occurs, we replace the LED board in the instrument and confirm the new board is within specifications. Another challenge has been the time required for instruments to cool down between runs, as the instruments do not include any active cooling. Refrigerated cooling blocks can be used to rapidly cool the block (2-3 mins), but this can be cumbersome at high throughput. Future improvements may include adding active cooling with a Peltier element.

Most other strategies for colorimetric LAMP do not allow for real-time readout with an accurate TTR. Although most diagnostic assays are qualitative, real-time readout is important for QC of these assays to ensure assay reagents react consistently within their specified readout time window. Realtime TTR can also enable quantitative measurements of target concentration. LARI provides a solution for real-time readout of TTR thanks to (a) rapid and repeatable conductive incubation of sample vials and (b) real-time monitoring of absorbance changes through each vial.

LARI has many potential applications beyond those presented here. Colorimetric LAMP, coupled with simplified, portable instrumentation, can be a powerful tool for global health and other field diagnostic applications. Additionally, the limited availability and throughput of conventional diagnostic instrumentation has presented a major bottleneck in responding to the COVID-19 pandemic. Simplified instrumentation such as LARI enables a robust, rapidly deployable alternative. Compared to conventional PCR thermocyclers, LARI requires relatively few low-cost components and can be assembled easily. This is particularly relevant in a pandemic situation when global supply chains are disrupted. We hope that the work presented here will aid the development of simplified diagnostic instrumentation and accelerate the broader adoption of colorimetric assays.

## Acknowledgements

The authors would like to acknowledge the help of Noor Alnabelseya, Dawn Spelke, Sangeeta Sarkar, and Sean Chang for experimental guidance and execution, Clay Reber for design consultation, James Provins for instrument assembly, and Fathom Manufacturing for design and fabrication of the urethane-cast housing.

